## Supplementary Information for "PMCE: efficient inference of expressive models of cancer evolution with high prognostic power"

### References

N. Simon, J. Friedman, T. Hastie, R. Tibshirani, Regularization paths for cox's proportional hazards model via coordinate descent, Journal of statistical software 39 (2011) 1.

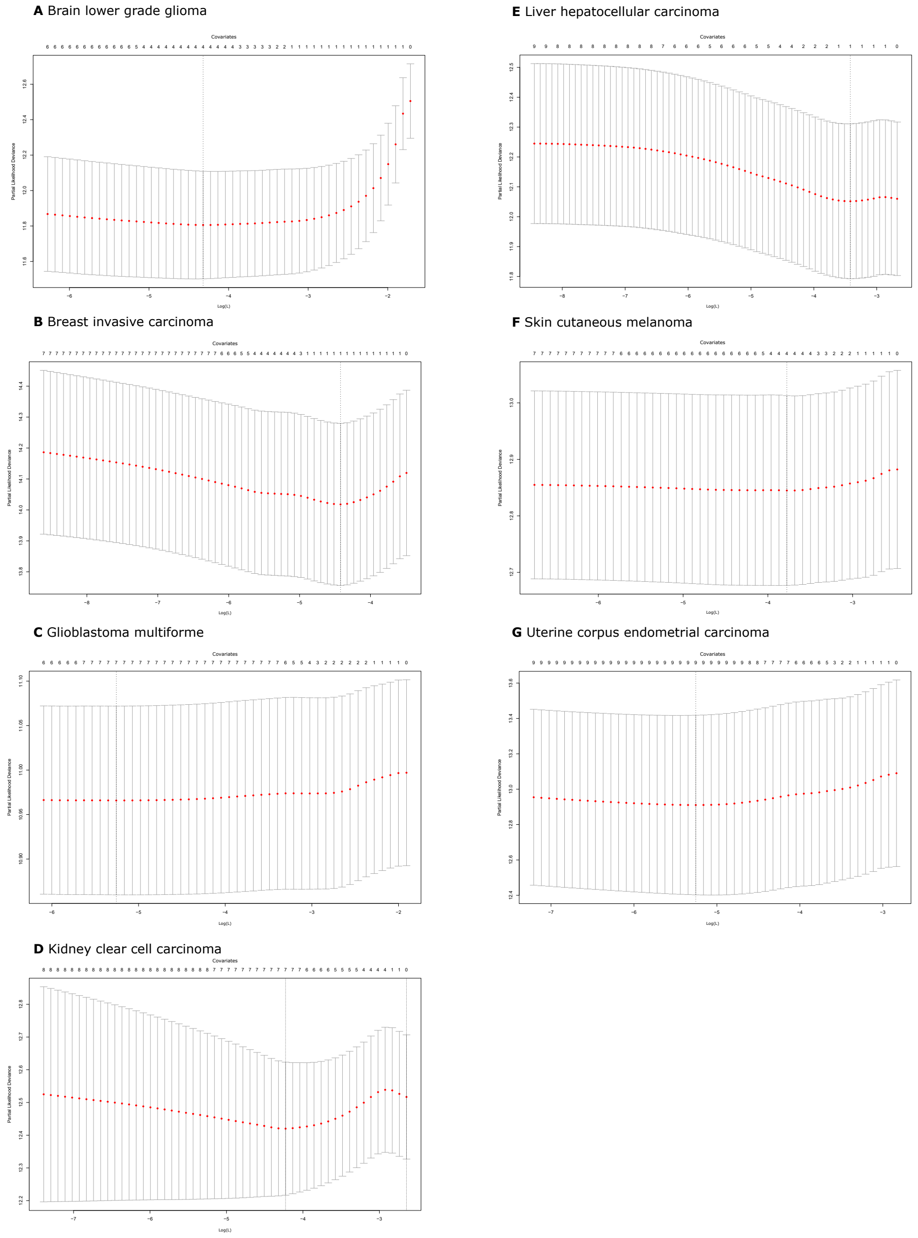


**Fig. 1.** Cross-validated error rates obtained with the method by Simon et al. (2011). The X axis is associated to the values of $L$ (i.e. the coefficient of the elastic net penalty), with error bars providing a confidence interval for the cross-validated error rate. The vertical bars indicate the minimum error. The X axis gives the optimal number of covariates for a model. In this figure, we present only the cancer types that show the size of the model different from 0.

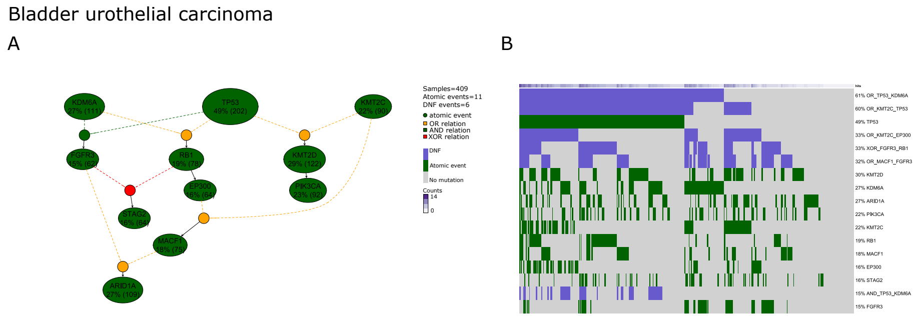


**Fig. 2.** A) PMCE model for bladder urothelial carcinoma; we show the atomic events (i.e. driver mutations on genes), the inferred formulas (AND, OR or XOR) and their prevalence in the considered cancer samples. B) Oncoprint of the input dataset for this tumor, where we report presence or absence of a each mutation for each sample.

Brain lower grade glioma

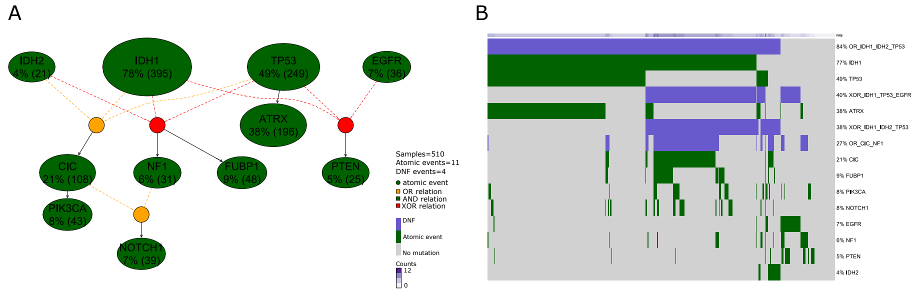


**Fig. 3.** A) PMCE model for brain lower grade glioma; we show the atomic events (i.e. driver mutations on genes), the inferred formulas (AND, OR or XOR) and their prevalence in the considered cancer samples. B) Oncoprint of the input dataset for this tumor, where we report presence or absence of a each mutation for each sample.

Breast invasive carcinoma

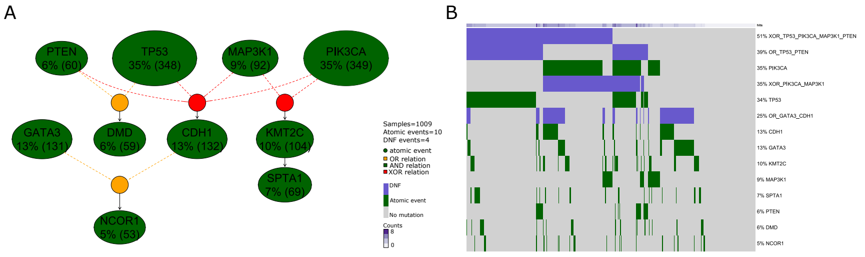


**Fig. 4.** A) PMCE model for breast invasive carcinoma; we show the atomic events (i.e. driver mutations on genes), the inferred formulas (AND, OR or XOR) and their prevalence in the considered cancer samples. B) Oncoprint of the input dataset for this tumor, where we report presence or absence of a each mutation for each sample.

Colorectal adenocarcinoma

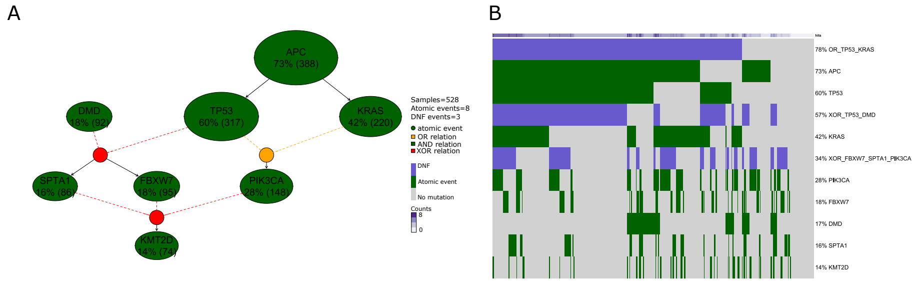


**Fig. 5.** A) PMCE model for colorectal adenocarcinoma; we show the atomic events (i.e. driver mutations on genes), the inferred formulas (AND, OR or XOR) and their prevalence in the considered cancer samples. B) Oncoprint of the input dataset for this tumor, where we report presence or absence of a each mutation for each sample.

Glioblastoma multiforme

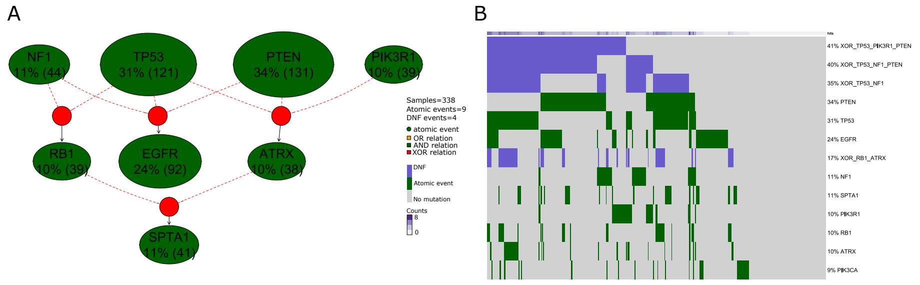


**Fig. 6.** A) PMCE model for glioblastoma multiforme; we show the atomic events (i.e. driver mutations on genes), the inferred formulas (AND, OR or XOR) and their prevalence in the considered cancer samples. B) Oncoprint of the input dataset for this tumor, where we report presence or absence of a each mutation for each sample.

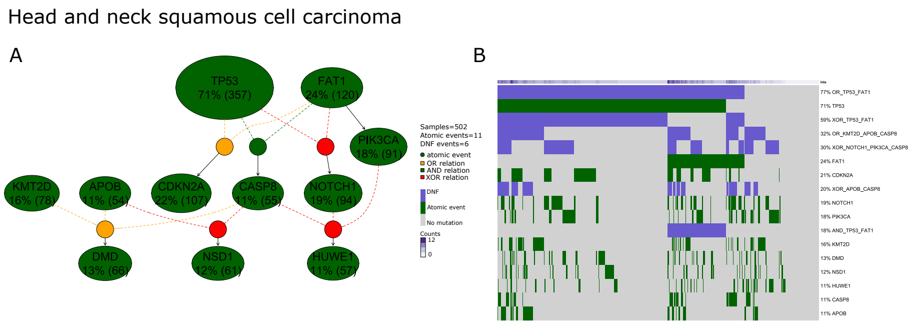


**Fig. 7.** A) PMCE model for head and neck squamous cell carcinoma; we show the atomic events (i.e. driver mutations on genes), the inferred formulas (AND, OR or XOR) and their prevalence in the considered cancer samples. B) Oncoprint of the input dataset for this tumor, where we report presence or absence of a each mutation for each sample.

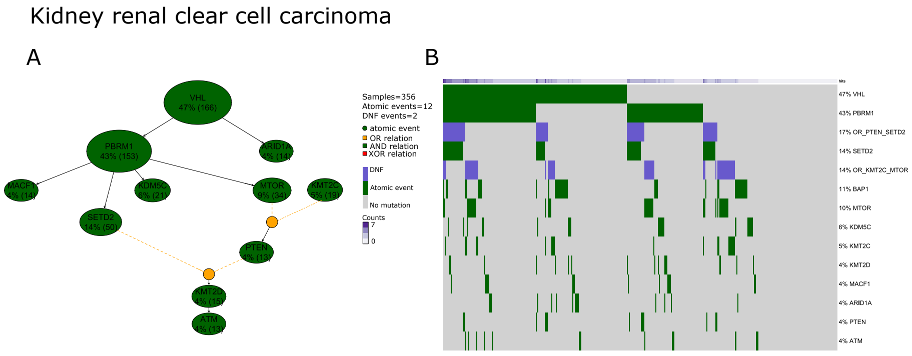


**Fig. 8.** A) PMCE model for kidney renal clear cell carcinoma; we show the atomic events (i.e. driver mutations on genes), the inferred formulas (AND, OR or XOR) and their prevalence in the considered cancer samples. B) Oncoprint of the input dataset for this tumor, where we report presence or absence of a each mutation for each sample.

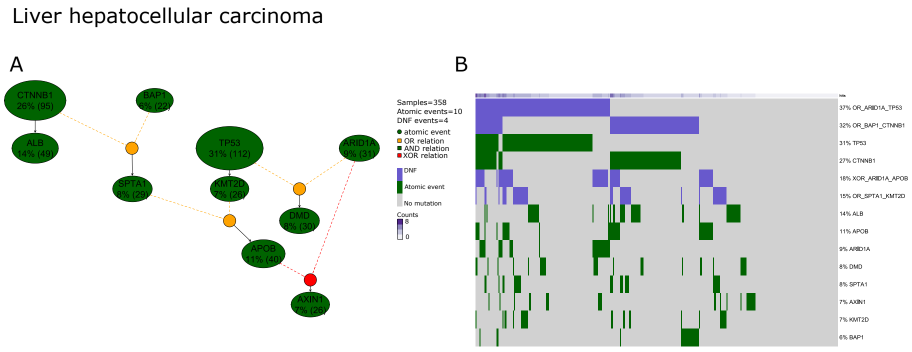


**Fig. 9.** A) PMCE model for liver hepatocellular carcinoma; we show the atomic events (i.e. driver mutations on genes), the inferred formulas (AND, OR or XOR) and their prevalence in the considered cancer samples. B) Oncoprint of the input dataset for this tumor, where we report presence or absence of a each mutation for each sample.

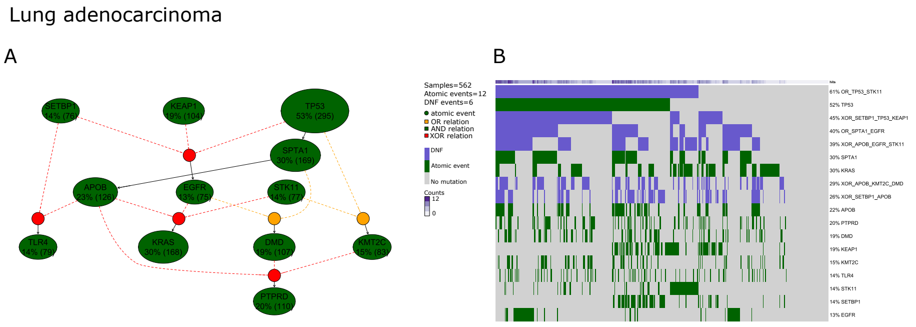


**Fig. 10.** A) PMCE model for lung adenocarcinoma; we show the atomic events (i.e. driver mutations on genes), the inferred formulas (AND, OR or XOR) and their prevalence in the considered cancer samples. B) Oncoprint of the input dataset for this tumor, where we report presence or absence of a each mutation for each sample.

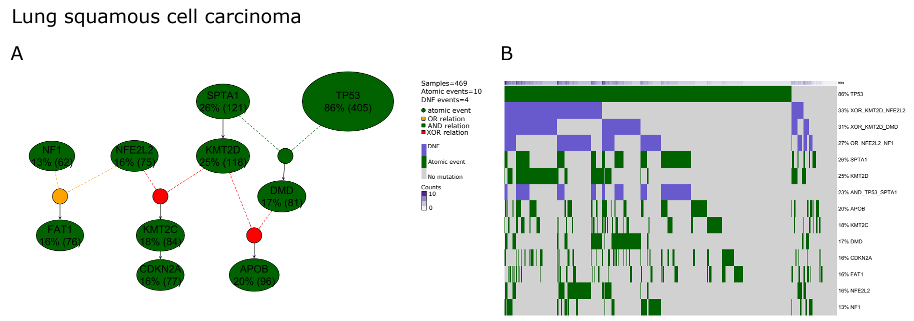


**Fig. 11.** A) PMCE model for lung squamous cell carcinoma; we show the atomic events (i.e. driver mutations on genes), the inferred formulas (AND, OR or XOR) and their prevalence in the considered cancer samples. B) Oncoprint of the input dataset for this tumor, where we report presence or absence of a each mutation for each sample.

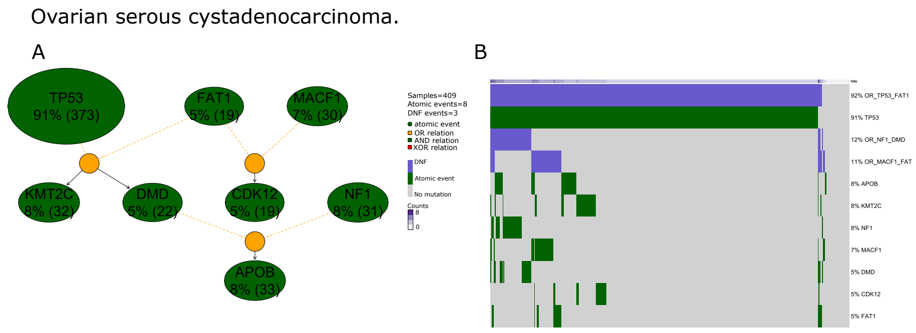


**Fig. 12.** A) PMCE model for ovarian serous cystadenocarcinoma; we show the atomic events (i.e. driver mutations on genes), the inferred formulas (AND, OR or XOR) and their prevalence in the considered cancer samples. B) Oncoprint of the input dataset for this tumor, where we report presence or absence of a each mutation for each sample.

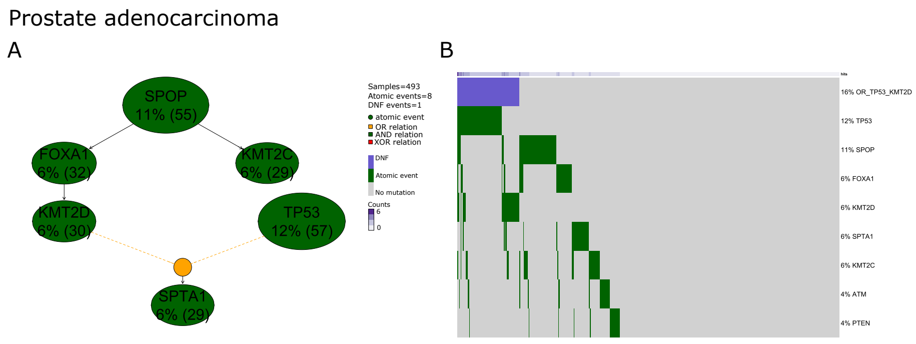


**Fig. 13.** A) PMCE model for prostate adenocarcinoma; we show the atomic events (i.e. driver mutations on genes), the inferred formulas (AND, OR or XOR) and their prevalence in the considered cancer samples. B) Oncoprint of the input dataset for this tumor, where we report presence or absence of a each mutation for each sample.

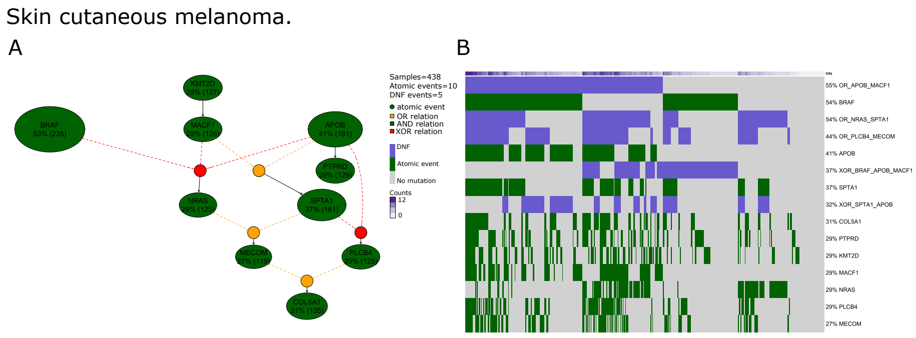


**Fig. 14.** A) PMCE model for skin cutaneous melanoma; we show the atomic events (i.e. driver mutations on genes), the inferred formulas (AND, OR or XOR) and their prevalence in the considered cancer samples. B) Oncoprint of the input dataset for this tumor, where we report presence or absence of a each mutation for each sample.

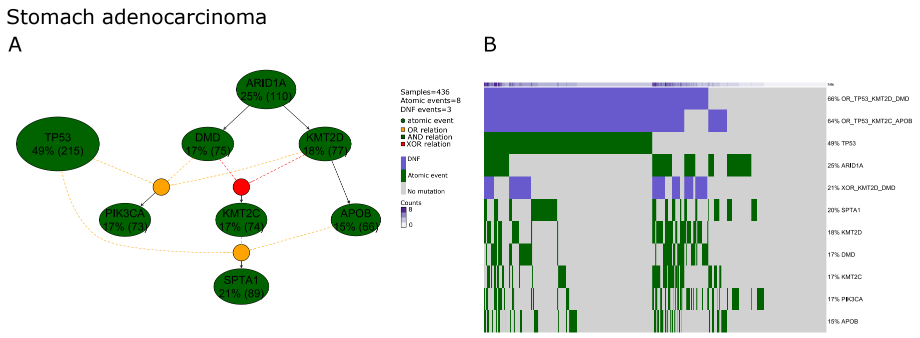


**Fig. 15.** A) PMCE model for stomach adenocarcinoma; we show the atomic events (i.e. driver mutations on genes), the inferred formulas (AND, OR or XOR) and their prevalence in the considered cancer samples. B) Oncoprint of the input dataset for this tumor, where we report presence or absence of a each mutation for each sample.

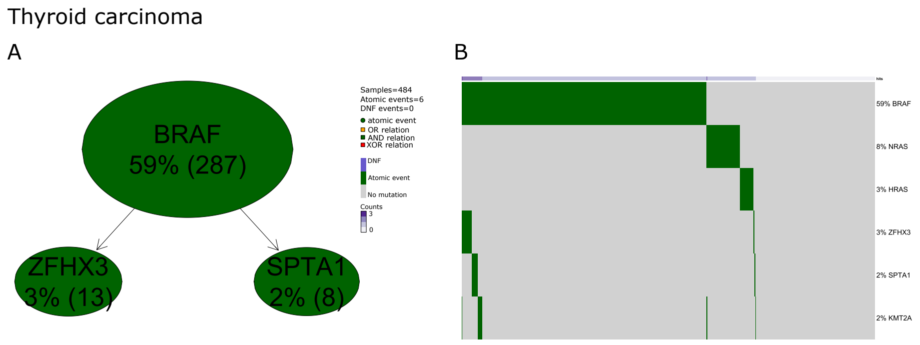


**Fig. 16.** A) PMCE model for thyroid carcinoma; we show the atomic events (i.e. driver mutations on genes), the inferred formulas (AND, OR or XOR) and their prevalence in the considered cancer samples. B) Oncoprint of the input dataset for this tumor, where we report presence or absence of a each mutation for each sample.

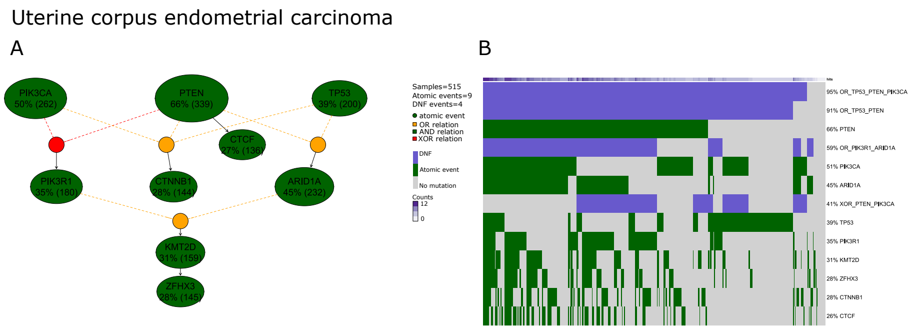


**Fig. 17.** A) PMCE model for uterine corpus endometrial carcinoma; we show the atomic events (i.e. driver mutations on genes), the inferred formulas (AND, OR or XOR) and their prevalence in the considered cancer samples. B) Oncoprint of the input dataset for this tumor, where we report presence or absence of a each mutation for each sample.

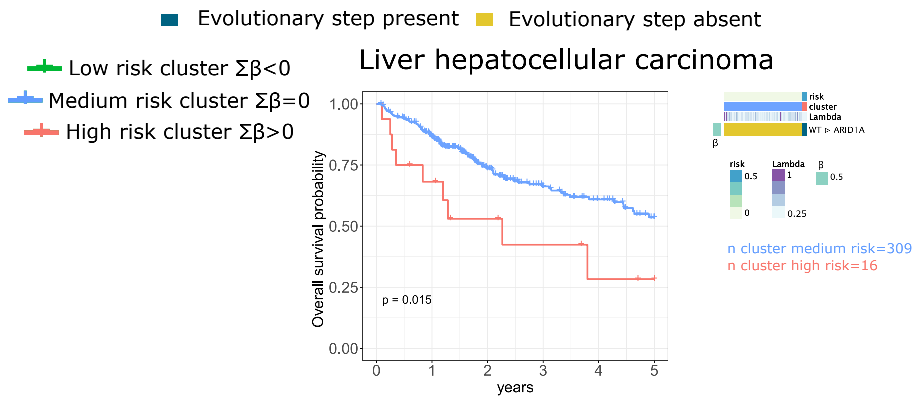


**Fig. 18.** Risk groups for Liver hepatocellular carcinoma.

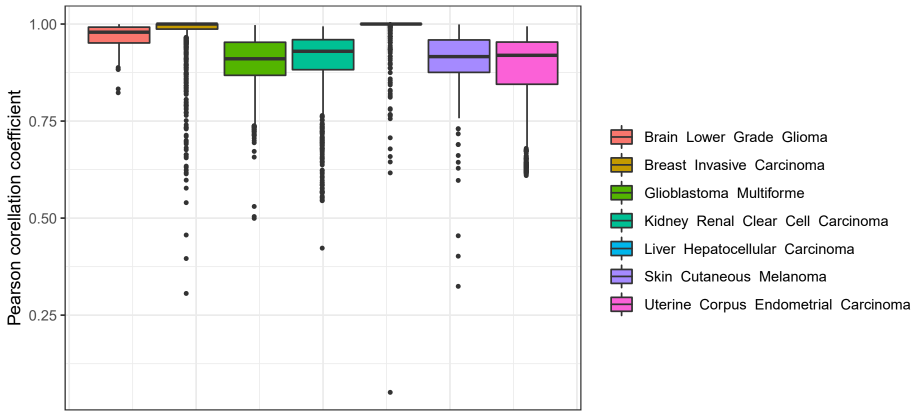


**Fig. 19.** Results of the cross-validation performed to estimate the stability of the survival analysis discussed in the main text. For each of the 7 cancer types that display a non-empty set of covariates associated to the minimum cross-validation error of the Regularized Cox Regression via Coxnet: $(i)$ 80% of the samples are selected, $(ii)$ the regularized Cox regression analysis is performed on the sampled dataset, $(iii)$ the sampling procedure is repeated 1000 times; $(iv)$ the Pearson correlation coefficient among the $\beta$ coefficients obtained from the sampled dataset and those from the full dataset is computed. The plot returns the distribution of the Pearson correlation coefficients for the 1000 independent sampling, with respect to each cancer type.

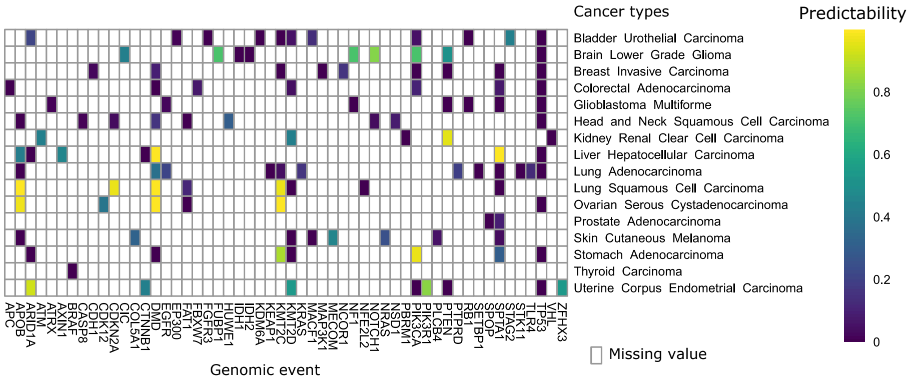


**Fig. 20.** Heatmap reporting the predictability score (range $[0, 1]$) for each genomic event included in the HESBCN models of the 16 considered cancer types. For each cancer type, for each genomic event, the predictability value of the subgraph defined by considering only the paths from the root to that event is computed as per Eq. (12) of the main text and reported. White cells in the heatmap indicates missing information, i.e. genomic events that are not present in the HESBCN of a given cancer type.

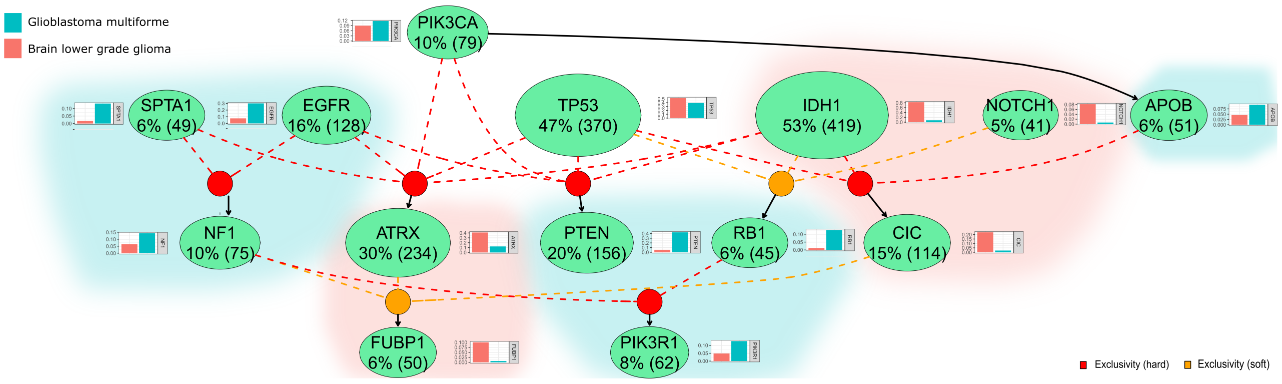


**Fig. 21.** HESBCN model for a pan-gliomas dataset including 510 lower grade gliomas (red) and 338 glioblastomas (green). Next to each genomic event, we show the barplot reporting the proportion of samples of the two distinct tumor types displaying that event; the molecular trajectories in which the prevalence of a given tumor is dominant are highlighted with a colored shades.

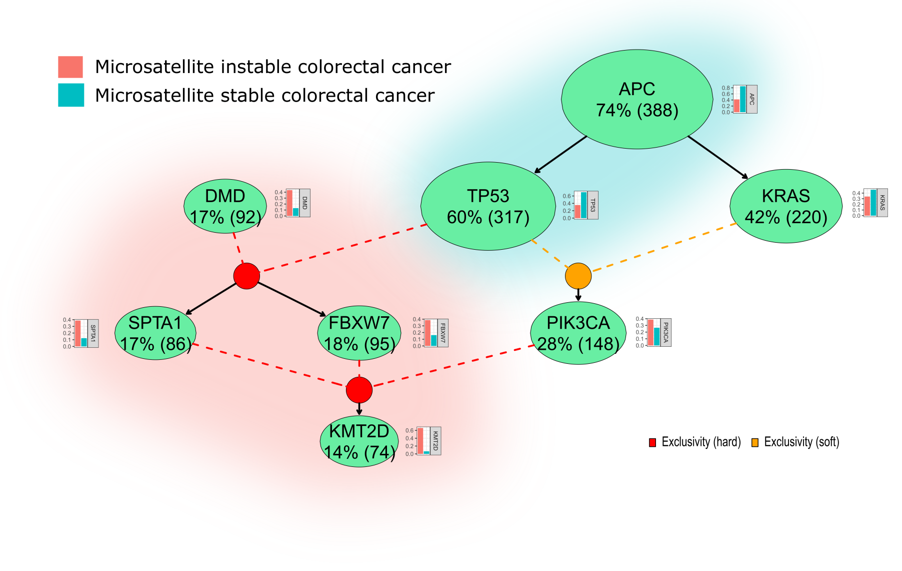


**Fig. 22.** HESBCN model for a dataset including 458 colorectal tumours comprising 396 microsatellite stable (MSS, green) and 62 microsatellite instable (MSI, red) tumours. Next to each genomic event, we show the barplot reporting the proportion of samples of the two distinct tumor types displaying that event; the molecular trajectories in which the prevalence of a given tumor is dominant are highlighted with colored shades.
